# Early-stage determinants of T1-S1 conformations in Kv1.3 channels

**DOI:** 10.1101/2025.07.09.663912

**Authors:** Liwei Tu, Aaron Sykes, Therese Davis, Sophia Gross, Carol Deutsch

## Abstract

Early-stage biogenesis of voltage-gated potassium channels (Kv) remains remarkably understudied yet is key to defining co- and post-translational acquisition of Kv secondary, tertiary, and quaternary conformations. Thus, we have studied nascent folding events and their determinants in Kv1.3, a tetrameric ion channel highly expressed in nerve and immune cells. We explore how folding of T1, an intersubunit recognition domain that ensures correct isoform assembly, is modulated by molecular determinants in the Kv subunit. Specifically, we focus on the T1-S1 linker and its highly conserved C-terminal sequence, S0. Using pegylation, a mass-tagging strategy, inter- and intrasubunit crosslinking, we define the molecular determinants of T1-S1 linker accessibility and location. Our findings show that i) dynamic protein-lipid and protein-protein linker interfaces exist, ii) the presence of a T1 domain and its conformation (monomer versus tetramer) impact linker properties, iii) helical formation of S0 occurs in early biogenic stages and does not require the presence of membranes, iv) a core recognition domain (T2) increases T1 dimerization efficiency and promotes tetramer formation, and v) as few as 12 native linker residues enable T1 tetramerization. These findings differ from canonical models and suggest that domain plasticity for Kv1.3 contributes to biogenesis and may underlie domain-domain communication in Kv function.

## INTRODUCTION

How does an ion channel form? It is a complex sequence of folding events, which happen early in biosynthesis of the channel protein and can ultimately affect the structure, function, and destination of the mature ion channel^1,2^. Misfolding due to a single mutation can cause pathology or even be lethal^3-5^. For voltage-gated K+ channels (Kv), we have made some progress in defining these early biogenic steps^6-14^ and find that cotranslational acquisition of local helices and turns begins in the ribosome exit tunnel and the ER membrane. These early events of secondary, tertiary, and quaternary folding are coupled.

Each subunit of a tetrameric Kv channel contains six transmembrane segments (S1-S6) consisting of a voltage-sensing domain (S1-S4) and a pore domain (S5, pore helix, selectivity filter, and S6) used for ion permeation. The cytoplasmic N-terminus of Kv1-4 families contains a highly conserved sequence (∼100 amino acids), the “T1” domain (Fig. 1). T1 serves as an intersubunit recognition domain that ensures correct isoform assembly^15,16^, kinetically facilitates subunit multimerization^17^, docks functionally important auxiliary proteins^18- 22^, underlies axonal targeting^23^, and undergoes voltage-dependent gating rearrangements at the T1-T1 interface^24-27^.

**Figure 1.**
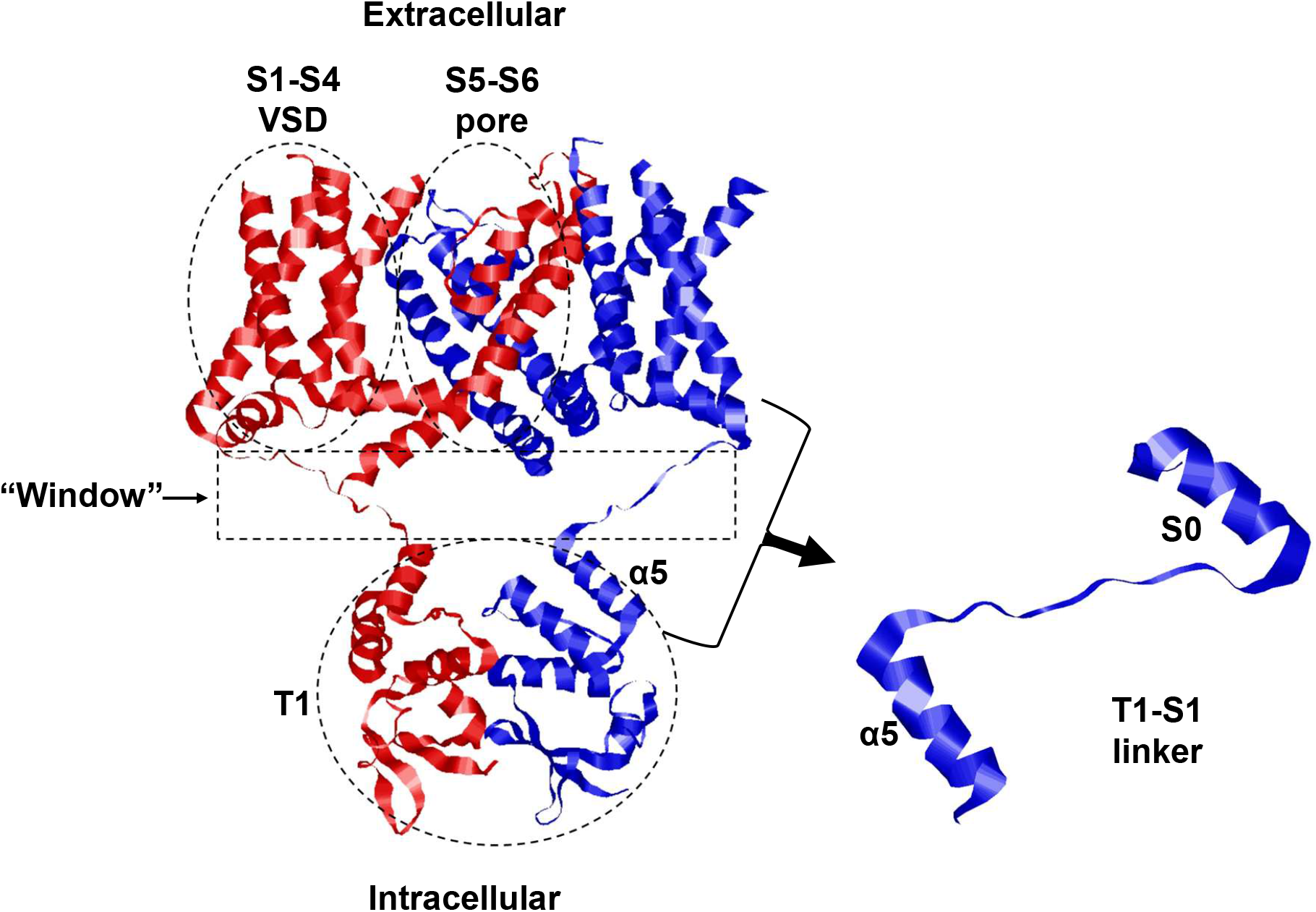
Kv 1.3 channel. Two adjacent subunits of the tetrameric mature Kv1.3 (PDB 7SSX) (left.) The assembled voltage-sensing domain (VSD, S1-S4), the permeation pore (S5-S6), cytosolic tetramerization domain (T1), the T1-S1 linker, and the “window” between T1 and the transmembrane segments (S1-S6) are highlighted. One subunit is colored red, one blue. On the right, a magnification of the T1-S1 linker in one subunit, including the S0 region shown as an α-helix reported in X-ray crystal and cryo-EM structures, and α5, which is the C-terminal segment of the T1 domain.

Here, we investigate a *Shaker*-like channel, human Kv1.3, which is highly expressed in nerve and immune cells. In T-lymphocytes, Kv1.3 is essential for activation and proliferation^28-32^. Moreover, it is causally linked to chronic inflammatory disease and autoimmune disorders, metabolic disorders, attention deficit and has been implicated in altered insulin sensitivity, neoplastic behavior and malignancy, and modulation of apoptosis ^33-35^. The severity of many of these pathologies correlates with Kv1.3 expression at the plasma membrane.

T1 is equally important in early biogenic events, especially as we discovered that Kv1.3 T1 folds both within, and upon emergence from, the ribosome exit tunnel^6,8^. The T1-S1 linker (42 residues), connects the T1 domain to the first transmembrane segment, S1, and includes a C-terminal sequence, S0, that has been modeled as an *α- helix* in the *mature* crystal structure^36,37^. We now explore how early-stage folding of the T1 domain is influenced by the T1-S1 linker. What is the role of this linker during Kv1.3 biogenesis? Is it a random spacer or is it encoded to mediate critical events in tertiary and quaternary folding? Three experimental observations are provocative and pose mechanistic questions for Kv biogenesis. First, S0 is *extended* in the Kv *nascent* protein inside the ribosome exit tunnel and upon its immediate emergence^12,38^. If indeed a helical S0 structure exists in the mature channel, this implies that both tertiary and quaternary interactions facilitate helix formation. What are these interactions, i.e., when, where, and why does S0 form a helix, and for what purpose? Using AGADIR, a helix propensity algorithm that predicts the helicity for soluble proteins^39^, we can infer the helix propensity of the T1-S1 linker sequence. Two regions flanking the T1-S1 linker, namely alpha5 (α5) at the C-terminal end of T1 (residues 142-153, numbering according to Cai et al.^40^) and transmembrane segment S1 (Fig. 1), each compact helices inside the exit tunnel^38^, serve as positive helical controls. T1-S1 residues flanking S0 N-terminally and C-terminally (154-162 and 173-182, respectively) do not have any helix propensity, whereas the S0 region, defined by residues 163-172, has a helix propensity similar to that calculated for the α5 sequence. Second, deletion of residues between T1 and S1, including S0, impairs channel function^16,41^but when co-expressed with WT yields functional heteromers, raising the question of which linker regions are required to permit correct Kv folding and function. Third, complete folding of T1 requires S0 be completely translated^7^. Does this ensure a pre-requisite length, location, and conformation of the linker for proper tertiary and quaternary T1 folding? This study investigates whether the linker is important in Kv channel formation and whether a new core recognition domain, “T2”^42-45^, exists and facilitates intersubunit interactions and multimerization to form a Kv channel.

## RESULTS

### Determinants of accessibility/location of T1-S1 linker residues

In each subunit of the mature Kv1.3, a 42- residue linker joins the T1 domain to S1 (Figs. 1, 2A, T1-S1 linker), which together in the mature Kv channel has been modeled as a “hanging gondola”^46^. This model has dominated the field yet remains experimentally and structurally unverified, especially in early biogenic stages. It predicts a linker location that is completely surrounded by water and fully accessible. Xray-crystal and cryo-EM structures involve detergent solubilization, which have limitations due to the experimental methods used and may not accurately reflect the conformation and location of loop regions (e.g., T1-S1 linker residues). We cysteine-scanned the linker and used a solvent accessibility assay, i.e., mass-tagging by pegylation (*Fpeg*) of accessible residues^38,47^, to experimentally identify the determinants of accessibility and location of each linker residue in the nascent monomer linker. Mass-tagging a single cysteine with methoxy-polyethylene glycol maleimide (PEG-MAL, 5000 Da) produces a ≥ 10 kD shift of the nascent peptide on a protein gel. Figure 2B shows the solvent accessibility of residues 155C, 169C, and 178C for ER membrane-inserted full-length monomer. Singly pegylated protein is the higher molecular weight band labeled “1” whereas unpegylated protein resides at the lower molecular weight labeled band “0”. Residue 155 is mostly pegylated while residues 169 and 178 are mostly unpegylated.

**Figure 2.**
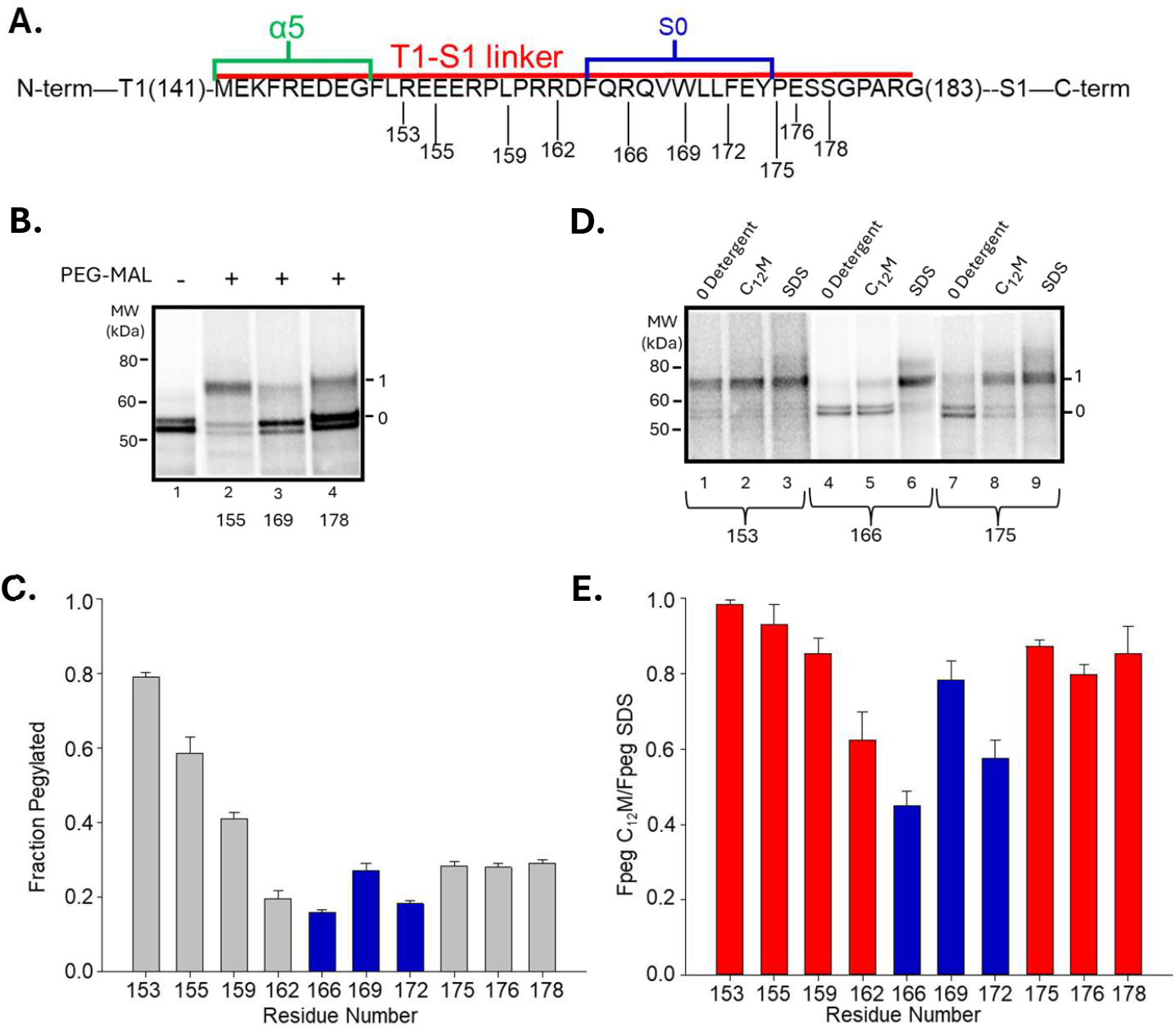
Accessibility and location of T1-S1 linker residues in monomeric Kv1.3. **A**. Kv1.3 amino acid sequence for the T1-S1 linker and α5. **B**. Solvent accessibility using pegylation of cysteine substitution at residues 155 (N-terminal T1-S1 linker), 169 (S0), and 178 (C-terminal T1-S1 linker) in membrane-inserted full-length Kv1.3. Unpegylated and singly-pegylated protein are marked as 0 and 1, respectively. The two intensities at band 0 are core glycosylated (upper) and unglycosylated (lower) protein. **C**. Fraction of each residue labeled with PEG-MAL (*Fpeg*) (n=3, mean ± SEM). Lower *Fpeg* values indicate less solvent accessibility. S0 bars are colored blue. **D**. Solvent-dependent relative accessibilities. Pegylation of residues 153, 166, and 175 in the absence of detergent, in C_12_M, or in SDS. Bands as defined in **B. E**. Relative ratio of *Fpeg* in C_12_M to *Fpeg* in SDS for T1-S1 linker residues (n=3, mean ± SEM). S0 bars are colored blue.

Surprisingly, a pegylation scan of the monomeric Kv1.3 linker reveals that most linker residues are *hindered* in the absence of detergents in early biogenesis (low *Fpeg*, Fig. 2C), specifically with increasing distance from T1. The most hindered (or least reactive) residues are in the S0 sequence and the C-terminal residues of the linker (*Fpeg* 0.15-0.30), suggesting they reside at lipid-protein and/or protein-protein interfaces during early biogenesis. To explore these hypotheses, we measured the ratio of *Fpeg* in C_12_M (non-ionic detergent) relative to *Fpeg* in SDS (anionic detergent) for each T1-S1 linker residue. C_12_M typically preserves the native state of a protein while disrupting protein-lipid and lipid-lipid interactions, but not protein-protein interactions. SDS typically unfolds and denatures proteins. All T1-S1 residues are maximally accessible in SDS (*Fpeg*∼ 0.8), an example of which is shown in Figure 2D for residues 153, 166, and 175 (lanes 3, 6, and 9, respectively). However, accessibility in zero detergent or C_12_M is residue-dependent (lanes 1vs 4, 2 vs 5, 5 vs 8) and indicates lipid-protein and/or protein-protein interactions. To assign these interactions, we calculated and compared *Fpeg* in each detergent. The normalized ratio of *Fpeg* in C_12_M to *Fpeg* in SDS will indicate which low*-Fpeg* residues are hindered by lipid and which by protein. A ratio close to 1.0 means the residue is maximally exposed (accessible) in both detergents but buried at a protein-lipid interface in zero detergent. A ratio <<1 suggests the residue is hindered at a protein-protein interface in C_12_M. N-terminal linker residue 153 is maximally exposed in zero detergent, C_12_M, and SDS, i.e., with equal relative band 1 intensities (lanes 1-3, Fig.2D), yielding *Fpeg* C_12_M/SDS ratios close to 1.0 (Fig. 2E). Therefore, residue 153 exists at a protein-aqueous interface, whereas nearby N-terminal residues 155-159 exist at both protein-aqueous and protein-lipid interfaces (compare Figs. 2C and 2E). C-terminal linker residues (175-178), while buried in zero detergent (low *Fpeg*, Figs. 2B (lane 4), 2C), have C_12_M/SDS ratios close to 1.0 (Figs. 2D (lanes 8 and 9) and 2E). These data suggest that lipid interactions in the native biogenic state contribute, either directly or indirectly, to produce the observed limited accessibility of C-terminal residues. In contrast, residues 166 and 172 in the S0 region remain modestly hindered, with C_12_M/SDS ratios of ∼0.4 and 0.6, respectively, consistent with occupancy at protein-protein as well as protein-lipid interfaces in the native state. The intervening 169 residue favors a protein-lipid interaction. We propose that S0 may be involved in Kv ‘core’-mediated functional interactions, e.g., interactions with membrane segments and/or the intervening loop segments. These results dispute the classic hanging gondola model and leave a new model to be discovered.

Are there any other molecular determinants of Kv1.3 able to influence accessibility/location? First, we asked whether the presence or absence of a folded T1 domain modulates linker location. Second, does the oligomeric status of T1, i.e., monomer versus tetramer, influence accessibility/location? To address the first case, we generated a full-length Kv1.3 that lacked the T1 sequence (T1-) by initiating translation at residue M142, followed by comparing *Fpeg* for linker residues in T1- and in full-length WT monomers. The accessibility of residues 159-172 in T1- monomer show a marked increase in C_12_M/SDS *Fpeg* (Fig. 3A), indicating a change in the molecular environment in this region. This suggests that the presence of the T1 domain in a WT monomer promotes a conformation in which S0 residues are more hindered, presumably by protein-protein interactions. Interestingly, most of the S0 residues in the T1- monomer lack protein-protein interactions. To address the second question of whether T1 tetramerization modulates linker accessibility, we compared pegylation of linker residues for the tetrameric and monomeric proteins (Fig. 3B). For this study, we used conditions that promote tetramerization (+membranes, 30°C, 2 hour-translation^6^) or monomer maintenance (±membranes, 22°C, 1 hour). *Fpeg* for N-terminal residues of the linker (155, 159, in zero detergent) increase in the tetramer, consistent with a more exposed conformation. These results suggest that T1 tetramerization relocates N-terminal linker residues further away from membrane lipid, potentially displacing the assembled T1 tetramer and expanding the ‘window’ above the T1 structure (Fig. 1). Such rearrangements may facilitate access and entry of ions, blockers, and peptide segments (e.g., an inactivation particle) to the permeation pore. Different locations of T1, mediated by linker rearrangements, sets a precedent for dynamic T1 conformations induced by functional states of the channel, e.g., during gating^25,27^. In contrast, S0 residues do not show any significant change in accessibility. Protein-protein and protein-lipid interactions with S0 may establish a more stable local S0 environment compared to N-terminal linker residues.

**Figure 3.**
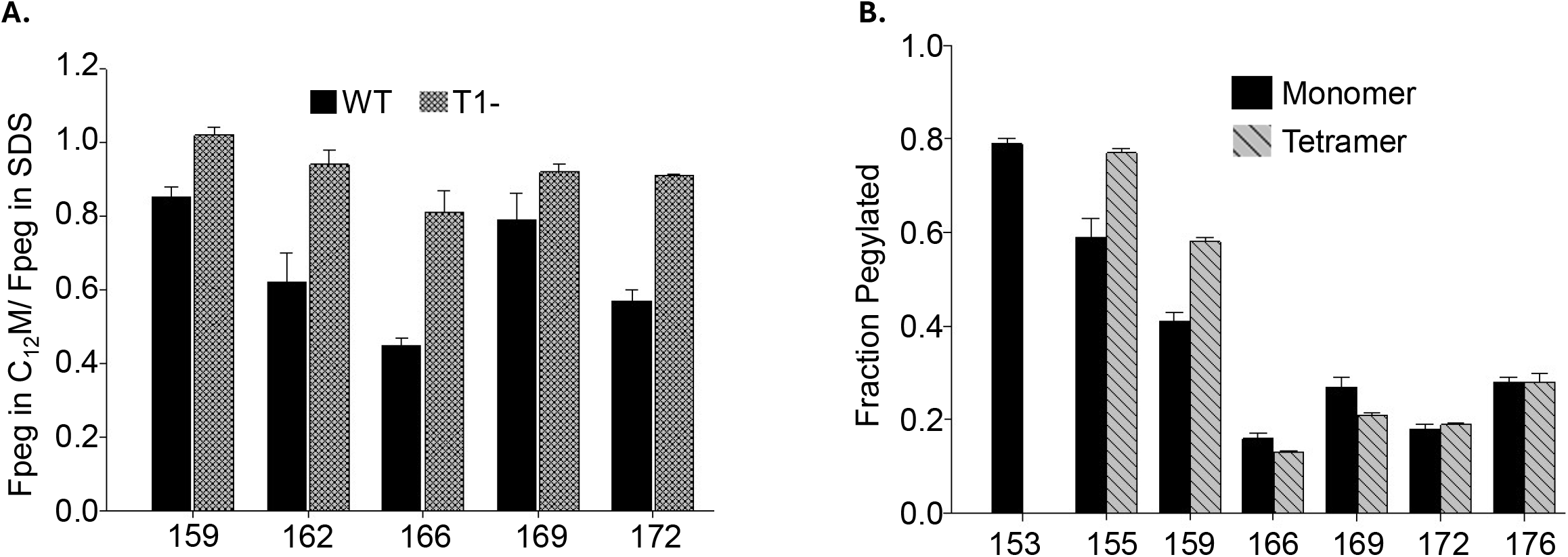
Modulation of linker location by the presence of T1 and T1-T1 Tetramerization. **A**. Solvent-dependent relative accessibilities for residues in the T1-S1 linker in both WT (solid bar) and T1– (stippled bar). For both constructs, translation was carried out at 22°C for one hour in the presence of membranes. Data represent means ± SEM, n=3. **B**. Solvent accessibility for T1-S1 linker residues in both monomer (solid bar) and tetramer (striped bar) (n=3, mean ± SEM). For the monomer, translation was carried out at 22°Cfor one hour in the presence of membranes. For the tetramer, translation was carried out at 30°C for two hours in the presence of membranes.

### Conformation of S0 during biogenesis

S0 contains a sequence, **FQRQVWLLF**, that is conserved across Kv1.x channels. It is a caveolin (Cav1) binding domain (CBD). Cav1-bound Kv1.3 targets to the plasma membrane, whereas impaired Cav1 binding leads to Kv1.3 intracellular retention, mitochondrial targeting, and facilitated apoptosis^48^. Multiple mutations of the CBD alter its binding affinity but do not inhibit tetramerization. However, electrophysiological properties (including steady-state activation and C-type inactivation) are altered^48^, implicating linker modulation of the Kv pore and/or voltage-sensor domains. While S0 has been described from X-ray crystal and cryo-EM structures as an α-helix in mature Kv tetramers^36,37^, the kinetics and determinants of its helix formation are unknown and provocatively important since we demonstrated that S0 is *fully* extended during its 100Å-passage through the ribosome exit tunnel and upon emergence^38^. Given S0’s biogenic and functional roles, it is important to further understand when and by what mechanism S0 forms an α-helix. This is unknown but will ultimately determine Cav1 binding and Kv1.3 localization.

Here we use an *intra*molecular crosslinking approach^13^ (*Pxlink*) to determine the conformation of S0 during biogenesis. An α-helix is defined as having 3.6 residues per turn and a pitch of 5.41Å per turn. Consequently, a pair of reactive and accessible cysteines engineered one, but not two or more, helical turns apart can be crosslinked with o-phenyldimaleimide (PDM), which has an intermaleimide distance of 6Å. A high probability of intramolecular crosslinking, *Pxlink*, is consistent with an α-helix. For Kv1.3, three pairs of cysteines, 165C/169C, 168C/172C, and 165C/173C, were separately engineered in S0. If S0 is an α-helix, then the first two pairs will each be separated by 5.41Å and crosslinked with PDM; the third pair will be >10Å apart and poorly crosslinkable with PDM. Our first goal was to further confirm that S0 forms a helix in tetrameric Kv1.3. We translated KpnI-cut mRNA that contained a cysteine pair, in the presence of membranes at 30° for 2 hours. These translation conditions promote membrane insertion and T1 tetramerization^6^. Intramolecular PDM crosslinking gave data that were analyzed using a maximum likelihood algorithm^13,49^ to simultaneously estimate *Pxlink* and the probability that available cysteine residues are labeled by PEG-MAL (*Ppeg*). *Ppeg*, for the indicated cysteines in all the aforementioned constructs was ∼ 0.7, enabling accurate calculation of *Pxlink*. Since residues in S0 are buried inside the membrane, we modified our crosslinking protocol. After 15 minutes of PDM incubation, C_12_M (1% final concentration) was added to promote the efficiency of *Pxlink* (see Methods/Discussion). Cysteine pairs 165C/169C and 168C/172C crosslinked to give *Pxlink* of 0.49±0.04 (n=3) and 0.48±0.04 (n=3), respectively, whereas a control, 165C/173C at a 2-turn distance, was not crosslinkable with PDM (Fig. 4B, right panel; Table 1). Two conclusions emerge from these results. First, our data are consistent with S0 helix formation in a mature Kv tetramer, thus confirming the proposed model from X-ray crystal and cryo-EM structures. Second, our findings indicate when and where S0 acquires a helical conformation, namely, early in Kv1.3 biogenesis prior to the complete translation of Kv1.3 C-terminal domains. To explore even earlier biogenic events and whether S0 helix formation occurs in the monomer and/or requires ER membranes and membrane insertion of the nascent Kv peptide, we determined intramolecular *Pxlink* in these same constructs in the absence of membranes, under conditions that favor monomer (22°C, 1 hour translation; Fig. 4B, left panel). *Pxlink* for 165C/169C and 168C/172C is 0.59±0.04 (n=5) and 0.58±0.04 (n=6), respectively. These findings suggest that S0, in the absence of membranes and membrane insertion, in a ribosome-attached nascent chain monomer, is a helix, even in the absence of T1 oligomerization. In summary, although S0 is fully extended inside the ribosome tunnel, it converts to a helical conformation as the nascent chain exits the ribosome and remains helical during biogenic tetramerization and in the mature Kv1.3.

**Table 1.** Accessibility and intramolecular crosslinking of S0. Cysteine pairs engineered in KpnI-cut Kv1.3 for intramolecular crosslinking assays of S0. The nascent peptides are attached to mRNA in the ribosome exit tunnel as described in Fig. 4. *Pxlink* is the probability that the cysteines are crosslinked and *Ppeg* is the efficiency of pegylation for the indicated cysteine pair. Data represent means ± SEM.

|  | Cysteine pairs | *Ppeg* | *Pxlink* |
| --- | --- | --- | --- |
| -mm<br>PDM (no C<sub>12</sub>M) | 165C/169C KpnI | 0.60 ± 0.01 (n=5) | 0.59 ± 0.04 (n=5) |
|  | 168C/172C KpnI | 0.65 ± 0.03 (n=6) | 0.58 ± 0.04 (n=6) |
|  | 165C/173C KpnI | 0.70 ± 0.02 (n=8) | 0.24 ± 0.02 (n=8) |
| +mm<br>1<sup>st</sup> PDM → 2<sup>nd</sup> C<sub>12</sub>M | 165C/169C KpnI | 0.67 ± 0.03 (n=3) | 0.49 ± 0.04 (n=3) |
|  | 168C/172C KpnI | 0.76 ± 0.01 (n=3) | 0.48 ± 0.04 (n=3) |
|  | 165C/173C KpnI | 0.66 ± 0.02 (n=3) | 0.28 ± 0.03 (n=3) |

**Figure 4.**
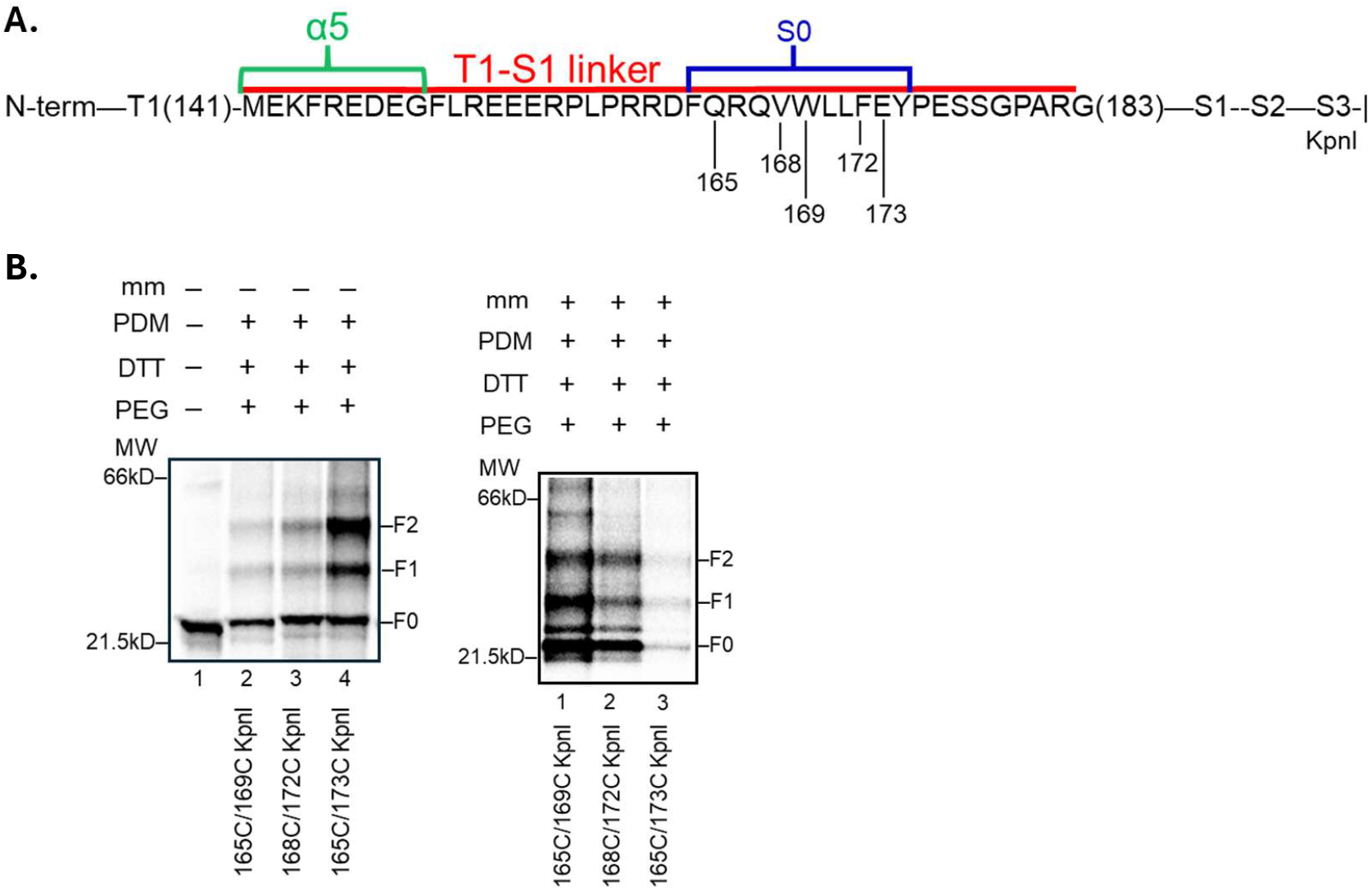
S0 conformation in nascent chain Kv1.3. **A**. Sequence of the T1-S1 linker as indicated in Fig. 2A including KpnI restriction site and S0 residues mutated to create engineered cysteine pairs 165C/169C, 168C/172C, and 165C/173C. **B**. Intramolecular crosslinking of ribosome-attached nascent peptides. Left gel: all lanes derived from translations of KpnI-cut mRNA in the absence of membranes. Lane 1 shows unmodified protein. Lanes 2-4 show crosslinking of 165C/169C, 168C/172C, and 165C/173C, respectively. Right gel: all lanes derived from membrane-containing translation of 165C/169C (lane 1), 168C/172C (lane 2), and 165C/173C (lane 3). Products were treated sequentially with PDM, DTT, and PEG-MAL. Symbols to the right represent doubly pegylated (F2), singly pegylated (F1) and unpegylated (F0) protein.

### Linker length-dependence for tetramerization

Having assessed the accessibility/location of T1-S1 linker residues and the specific conformation of S0, we asked whether the linker has a prerequisite end-to-end length and specific residue regions that are critical for tertiary and quaternary T1 folding. To address this question, we engineered a series of linker deletion constructs in a 118C/126C background (Table 2) and carried out two assays. First, we measured the fraction of peptide crosslinked as dimers, trimers, and tetramers (Fig. 5B, asterisks, and Fig. 5C). Second, we analyzed the fraction of crosslinked protein captured as trimers and tetramers, which reflects the efficiency of dimerization of dimers to form trimers and tetramers, the dominant pathway in the assembly of functional Kv1.3 tetramers^50^. Four deletion constructs with residues in the N-terminal region of the T1-S1 linker have significantly lower yields of assembled multimers, i.e. intermolecular *Pxlink* (bracketed constructs, Fig. 5C). Yet, of these four, only one deletion construct (Δ153-181) has a marked reduction in trimer/tetramer concentration (Fig. 5D). Decreased multimerization occurs for all four constructs, but (Δ153-181) additionally manifests impaired dimerization of dimers. All other deletion constructs are similar to WT, for which total protein crosslinked is ∼0.63 ± 0.01 (n = 43) and trimer/tetramer fractions are ∼ 0.84 ± 0.02 (n = 43). Deletion of the α5 segment, in the N-terminus of the linker (Δ142-152 and Δ143-172), or the highly charged REEER sequence (Δ153-157) do not disrupt multimer formation or efficiency of dimerization of dimers. Nor does deletion of the PLP motif in Δ153-160 or introduction of prolines into putative helices, namely α5 or S0 (D148P/E149P and R166P/Q167P, respectively) alter dimerization efficiency. The absence of S0 alone (Δ160-173) does not influence multimer formation, however in combination with other linker deletions (Δ153-181, Δ154-167), a marked decrease in the multimer fraction is observed (0.44 ± 0.05 and 0.32 ± 0.1, respectively, n = 3) with only Δ153-181 inhibiting dimerization of dimers (see above). These findings suggest that specific linker sequences contribute to the ability of the linker to promote T1 multimerization. This conclusion is supported by a comparison of Δ143-172 with Δ153-181, Δ153-157 with Δ157-160, Δ153-160 with Δ161-167, and Δ154-167 with Δ160-173. Each deletion pair has approximately the same number of remaining residues in the linker (Table 2) yet shows a significantly different probability of T1 association to form dimers, trimers, and tetramers (Fig. 5C). Moreover, as few as 12 linker amino acids can promote T1-T1 tetramerization. From these studies, we conclude that the T1-S1 linker is more than a simple random spacer of given end-to-end length. Rather, the linker has specific conformations conferred by its sequence, physicochemical interactions, and length, which impact T1 tetramerization, a fundamentally new role for the T1-S1 linker. We note, however, that there are some limits to interpretation of deletion studies. For example, deletions can produce amino acid proximities that perturb or promote new side-chain interactions, rotamerization of remaining residues, or conformational changes, some of which might impinge on multimerization. Despite these caveats, our results do provide insights that may bring us closer to understanding biogenic determinants of Kv tetramerization.

**Table 2.**
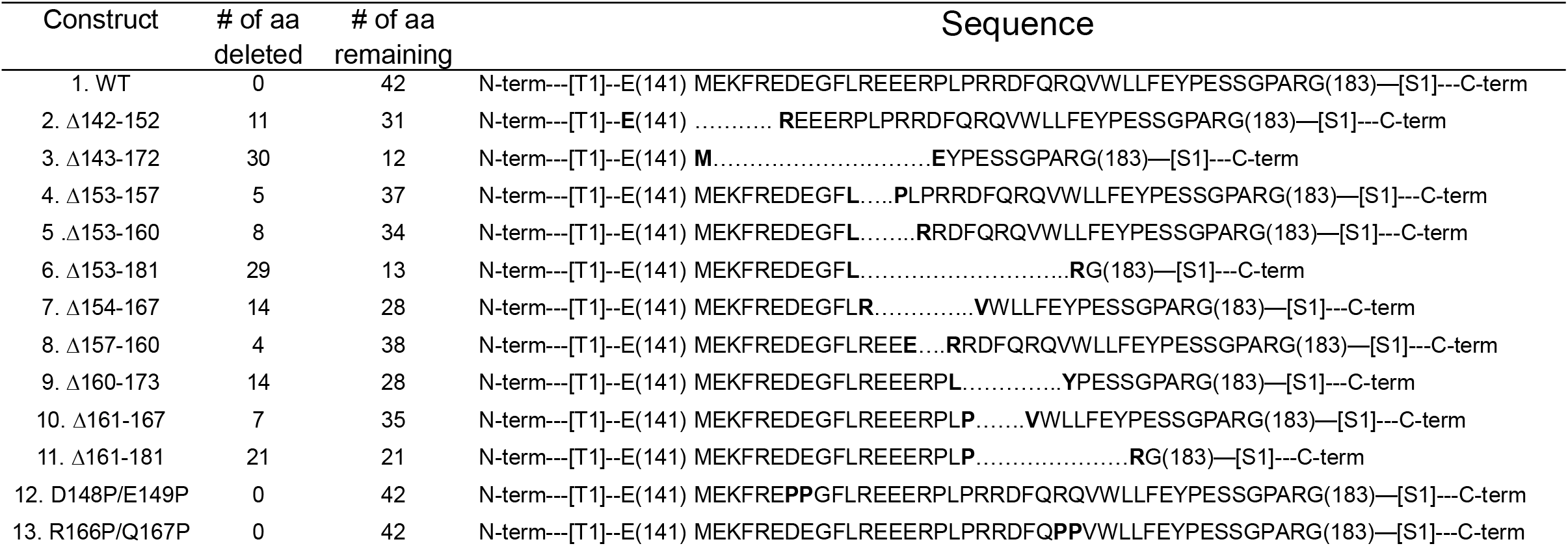
Deletion constructs and point mutations used in intermolecular crosslinking assays. The sequences shown are derived from the Kv1.3 T1-S1 linker M142-G183, which includes α5 (M142—E149), in a background that contains cysteines 118C/126C at the intermolecular T1-T1 interface.

**Figure 5.**
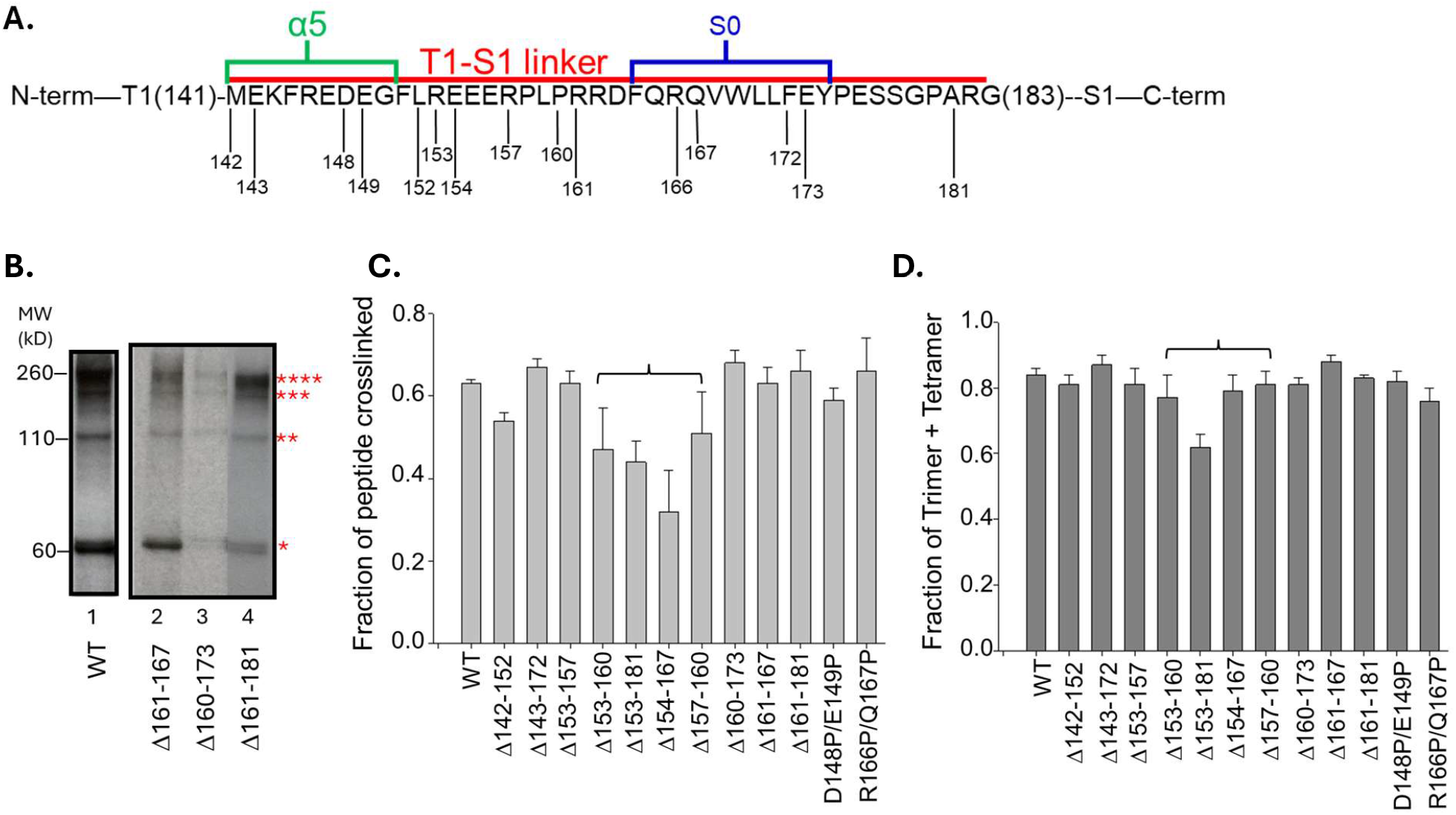
Specific regions of T1-S1 differentially affect tetramerization. **A**. Kv1.3 amino acid sequence as indicated in Fig. 2A, including residues used to generate ten deletion constructs and two mutation constructs (+PP). **B**. Quaternary folding of T1 domains. Multimer formation for WT (lane 1) and for T1-S1 deletion constructs (lanes 2–4). Modification with PDM yields monomer, dimer, trimer, and tetramer, indicated by the number of asterisks, respectively. Two cysteines in the T1 –T1 interface (118C/126C) were engineered in full-length EcoRI-cut background for this purpose. **C**. Fraction of total protein crosslinked as dimer, trimer, tetramer divided by the sum of monomer, dimer, trimer, and tetramer (mean ± SEM), i.e., equivalent to the probability of intermolecular crosslinking (*Pxlink*). **D**. Fraction of crosslinked protein that is trimer + tetramer, calculated as the sum of trimers + tetramers divided by the sum of dimers, trimers, and tetramers.

### T2 domain promotes biogenic dimerization of T1 dimers

Previously, we showed that Kv1.3 lacking its N-terminal T1 recognition domain can tetramerize to form functional channels (albeit with lower efficiency and higher [mRNA] required) and that ‘core’ Kv1.3 fragments can, by direct association, suppress channel formation of co-expressed full-length WT^43,51^. Both findings compellingly support the existence of a recognition domain (“T2”) in the core S1-S5, that contributes to oligomerization efficiency, consistent with findings in other *Shaker*-like Kv channels lacking T1 and part of the T1-S1 linker^17,25,42^. This represents a conceptual modification of current dogma. To explore whether T2 affects T1 multimerization, we created an enzyme-cut *KCNA3* DNA digested at the KpnI-site (Fig. 6A). In our translation system, KpnI-cut mRNA cotranslationally integrates into the membrane, gets core glycosylated in the S1-S2 loop, and produces T1 tetramerization^6,44^. There is no stop codon in KpnI-cut mRNA. Translation is stalled at the end of the S3 segment, leaving S3 attached to mRNA inside the ribosome exit tunnel whereas S1 and S2 translocate across the ER membrane. This construct is thus missing much of the Kv core region (putative T2 contributors). We translated this KpnI-cut mRNA and EcoRI-cut mRNA (full-length) in the presence of membranes for 2 hours. Intermolecular PDM crosslinking of both KpnI-cut and EcoRI-cut proteins gave monomer, dimer, trimer, and tetramer (asterisks, Fig. 6B) at translation temperatures of 30°C and 22° (2 hours), both of which enable tetramer formation. The full-length Kv1.3 and core-truncated Kv1.3 proteins have similar total *P*xlink, i.e., fraction of intermolecular crosslinked protein (data not shown). However, the truncated KpnI-cut fragment yields a dramatically reduced pool of trimers and tetramers with proportionately more dimers present (Fig. 6B and 6C), i.e., a redistribution of multimeric species that reflects protein stalled at the dimer stage. We propose an interfacial conformation of the KpnI-dimer is energetically unfavorable for T1 dimer-dimerization and requires the rest of the core Kv1.3 T2 to stabilize a productive dimeric conformation for tetramer formation, a process that increases the efficiency and kinetics of T1 dimerization of dimers^50^. Additionally, lower temperature partially recovers the trimer/tetramer yield, perhaps mediated by a permissive dimer conformation that facilitates dimerization of dimers. Our findings suggest a new model of biogenic Kv folding, one in which nascent chain elongation and availability of a complete T2 domain is required to efficiently shift the multimeric equilibrium toward T1 tetramers.

**Figure 6.**
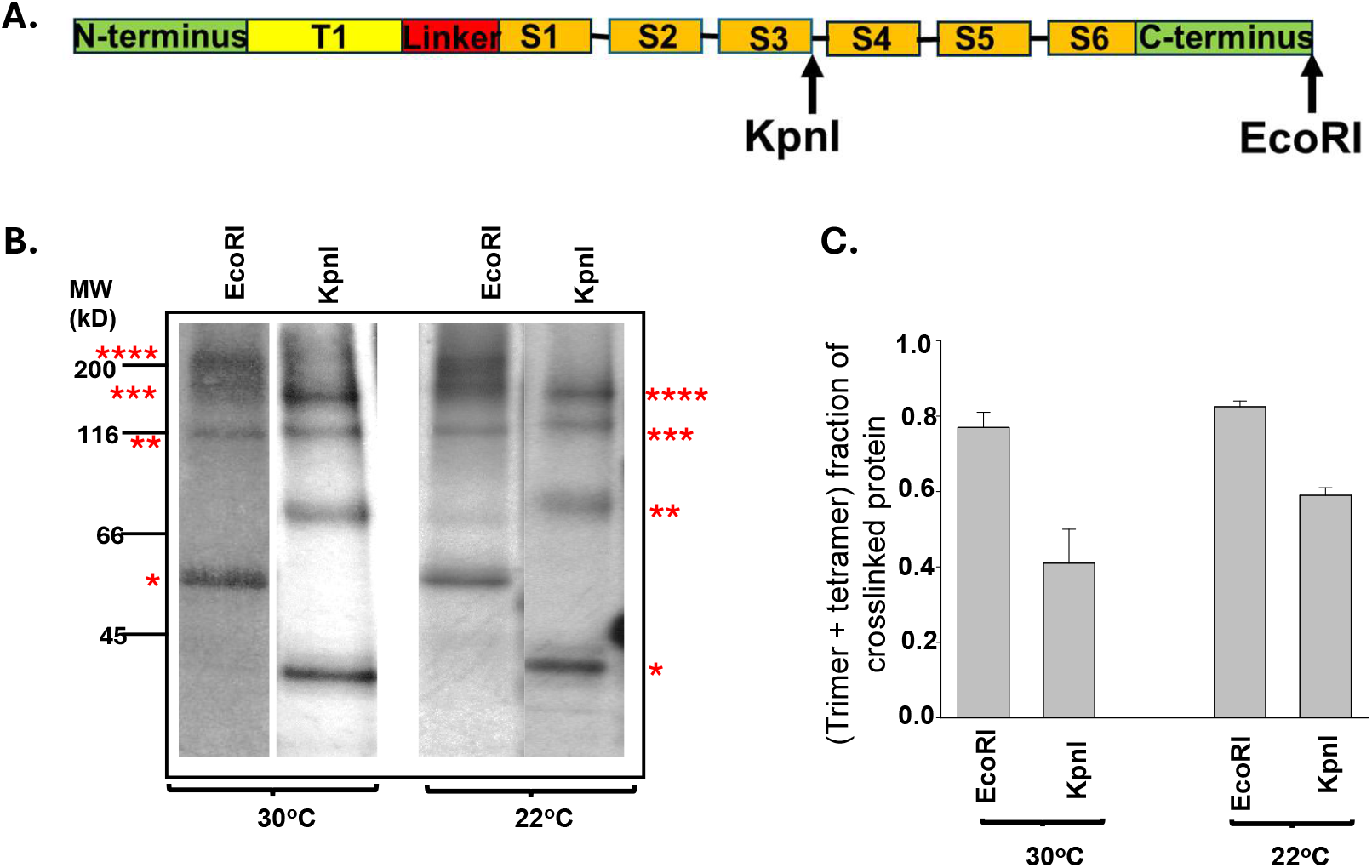
Core Kv segments lead to a redistribution of multimeric species. **A**. Sequence of Kv1.3 showing restriction sites. **B**. Intermolecular crosslinked multimers for the indicated cut-sites KpnI and EcoRI in a 118C/126C Kv1.3 background. Modification with PDM yields monomer, dimer, trimer, and tetramer, indicated by the number of asterisks, respectively. Translation was carried out at the indicated temperatures in the presence of microsomal membranes. **C**. Fraction of crosslinked protein present as trimer + tetramer as described in Fig. 5D.

## DISCUSSION

Our study explores the impact of the cytoplasmic T1-S1 linker of Kv1.3 on early biogenic folding and assembly of the T1 domain, a process that is highly optimized to promote efficient expression of the tetrameric channel. The T1-S1 linker connects the T1 domain to the transmembrane core of the channel but the impact of this relationship on folding and gating has been minimally investigated. Trypsin-cleavage of the T1-S1 linker is relatively slow compared with cleavage of an extracellular Kv loop control (S1-S2 loop), suggesting that the T1-S1 linker is not completely accessible, is partially structured, and/or contains dynamic structure^52^. Our pegylation studies now show this to be true. The T1-S1 linker is not a uniformly accessible loop and maintains conformations that place specific regions of the linker at lipid-protein and/or protein-protein interfaces. Both conclusions deviate from the hanging gondola model^46^. Furthermore, our findings suggest that the presence of T1 and/or its oligomeric state influence the location and conformation of the T1-S1 linker. Our measurements of linker accessibility reveal that N-terminal residues of the T1-S1 linker in the WT tetramer vis-à-vis the WT monomer are relatively more distant from the membrane, consistent with relocation of the linker (linear or angular distance) and/or rotamerization of the reporter cysteine. Both cases will expand the dimensions of the tetrameric “window” to accommodate occupancy of ions, peptides, and blockers. Alternatively, our accessibility results of S0 show no significant difference between WT monomer and tetramer, suggesting that the buried S0 may serve as a molecular anchor that modifies the window geometry and T1 location.

Although deletion of the T1 domain from several Kv isoforms still permits protein expression of functional channels, the efficiency is quite low^16,17,42,43,53,54^. In some cases, partial deletion of different regions of the T1- S1 linker abolishes formation of functional channels^41^, which is rescued by coexpression with full-length channel subunits. In our current studies, we find S0, a highly conserved sequence in the T1-S1 linker of the Kv1.x family, manifests protein-protein interactions in WT Kv1.3, but lacks this interaction in the T1- construct. Therefore, we propose that low Kv protein expression and altered gating produced by T1 deletions or mutations^17,25,42,43^ may be mediated by S0 interactions with the core transmembrane region, T2, in the assembled tetrameric channel. A similar inference may be drawn for voltage-sensor movements that induce T1 rearrangements^27^. This hypothesis is consistent with another finding that complete translation of T2 and its oligomerization increase T1 dimerization efficiency and promote tetramer formation. This may reflect a direct or indirect interaction between T1 and T2 and may involve S0.

Previous studies show that S0 is extended inside the tunnel^38^and is suggested to be an α-helix in structures of the mature Kv1.3 channel^37^. Our current findings describe the intervening events. First, S0 forms a helix upon emergence form the tunnel. Second, this occurs while the monomeric peptide is still attached to the ribosome, prior to complete translation of C-terminal Kv domains. Third, S0 helix formation is independent of membrane insertion of downstream transmembrane segments and T1 tetramerization. We find that S0 is helical in tetrameric Kv1.3 *(Pxlink* ∼ 0.5) in the presence of membranes. A slightly higher *Pxlink* of 0.6 was measured in the absence of membranes, consequent to relief of hindrance or relief of S0 conformational rearrangement in lipids. In contrast, after complete solubilization of the membrane prior to PDM addition, we were unable to detect bifunctional crosslinking (*Pxlink* 0.3, data not shown). This observation may further suggest that the lipids and/or local environment induce different S0 conformations, including helical rearrangements with relatively lower stability. Such induced S0 rearrangements may facilitate transmission of signals/force between T1 and Kv core domains during channel function. We may speculate why S0 helix formation occurs early in the biogenic process. Chaperone interactions or binding reactions (e.g., caveolin binding^48^ may be time-dependent for efficient folding, assembly, and targeting to functional compartments, namely mitochondria versus the plasma membrane. Later in the biogenic process, during multimerization, S0 may be buried in lipid.

The dominant pathway in tetramer formation is dimerization of dimers and the steady-state concentration of trimers is relatively low^50^. Moreover, dimerization likely uses interaction sites different from those involved in monomer-monomer association^50^. Our study of deletion mutants in the Kv1.3 T1-S1 linker revealed both length- and sequence-specific dependence for multimerization. Several deletions yielded a decrease in the fraction of total protein crosslinked as dimers, trimers, and tetramers. However, except for one deletion mutant (Δ153-181), the fraction of dimers in the deletion peptides is similar to that in WT. Consequently, we speculate that the initial association rate of T1 monomers in some deletion mutants is slower than that for WT, i.e., rate-limiting, and that subsequent dimerization of dimers is unaffected by T1-S1 linker deletions. The underlying reasons may be twofold. First, the newly formed dimer surface speeds kinetics of association with a proximal dimer. Second, the newly formed dimer surface creates a potent affinity for dimer-dimer association. A final intriguing speculation is that S1 likely has conformational plasticity. Helical interconversion of S1 (α ↔3_10_) and S1 tilting will influence multimerization efficiency, T1 location, and the potency of T2 interactions. For example, if the linker length is shortened by deletion, folding, or made inaccessible by association with lipid or protein, then a compensatory adjustment in S1 could enable as few as 12 linker residues to mediate tetramerization. In summary, we note that multimer association is subject to several governing determinants and may involve intrasubunit and intersubunit contributions by T1, the T1-S1 linker, and T2 domains.

## METHODS

### Constructs and in vitro translation

Standard methods of bacterial transformation, plasmid DNA preparation, and restriction enzyme analysis were used. The nucleotide sequences of all mutants were confirmed by automated cycle sequencing performed by the DNA Sequencing Facility at the University of Pennsylvania School of Medicine on an ABI 3730XL Sequencer using BigDye terminator chemistry (A0BI). Engineered cysteines and restriction enzyme sites were introduced into pSP/Kv1.3/cysteine-free using QuikChange Site-Directed Mutagenesis Kit (Agilent, Santa Clara, CA). Kv1.3 residue numbers begin at the 2^nd^ ATG site^40^. All mutant DNAs were sequenced throughout the region of the mutation. Capped cRNA was synthesized in vitro from linearized templates using SP6 RNA polymerase (Promega, Madison, WI). Linearized templates for Kv1.3 translocation intermediates were generated using several restriction enzymes to produce different length DNA constructs lacking a stop codon and to position the nascent peptide segments at the specified locations. Proteins were translated in vitro with [^35^S]methionine (2 μl/25 μl translation mixture; ∼10 μCi/μl Express (EasyTag), Revvity Health, Boston, MA) for 1 hr at 22°C or 2 hr at 30°C in a rabbit reticulocyte lysate (Promega, Madison WI; 2 mM final [DTT]) with 1 μl of microsomal membranes (canine pancreas: gift from Gunnar von Heijne; Promega, Madison WI) when applicable according to the Promega Protocol and Application Guide.

### Pegylation

Translation product (25 μl) was diluted into PBS containing 2 mM DTT and centrifuged through a sucrose cushion (120 μl; 0.5 M sucrose, 100 mM KCl, 5 mM MgCl_2_, 50 mM HEPES,1 mM DTT, pH 7.3) for 7 min at 50,000 rpm or 22 min at 70,000 rpm with a TLA 100.3 Beckman ultra-centrifuge rotor at 4 °C to isolate membrane-bound or ribosome-bound peptide, respectively. The supernatant was completely removed, and the pellet was resuspended on ice in 50 μl of PBS containing 1 mM DTT with either 1% dodecyl maltoside (C_12_M), 2% SDS, or no detergent and incubated for 1 hour. Samples suspended in the absence of detergent or in C_12_M were incubated on ice while SDS samples incubated at room temperature. Pegylation was started by adding 50 μl of 10 mM polyethylene glycol maleimide (PEG-MAL, MW 5000; SunBio Korea) to give a final concentration of 5 mM PEG-MAL and incubated for 2 hours on ice. The reaction was then terminated by adding 100 mM DTT and incubating at room temperature for 15 minutes. After pegylation, samples were precipitated with acetone containing 0.4 mM HCl overnight. Pellets were dissolved in 14.5 μl DPBS buffer, 2.5 μl of DTT (10×), and 7 μl of LDS (4×).

### Intermolecular Crosslinking Assay

Translation reaction (25 μl) in the presence of microsomal membranes (1 μl) was added to 500 μl of PBS buffer, followed by incubation with 0.5 mM ortho-phenyldimaleimide (o-PDM) for 60 min on ice. The reaction was terminated by adding 5 mM DTT and incubated at room temperature for 15 min. Then, the reaction was centrifuged through a sucrose cushion (120 μl; 0.5 M sucrose, 100 mM KCl, 5 mM MgCl_2_, 50 mM HEPES (pH 7.3) and 1mM DTT) at 4°C for 7 min at 50,000 rpm using a TLA 100.3 Beckman Optima TLX Ultracentrifuge, to isolate membrane-bound peptide. The supernatant was aspirated, and the pellets were dissolved in 14.5 μl DPBS buffer, 2.5 μl of DTT (10×), and 7 μl of LDS (4×).

### Intramolecular Crosslinking Assay

Translation reactions (25 μl) were carried out in rabbit reticulocyte lysate, 2 mM DTT (Promega Protocol and Application Guide) both in the presence and absence of canine microsomal membranes (1 μl) using KpnI-cut cysteine-free constructs containing engineered cysteines. A 12 μl sample without membranes was diluted into 700 μl of 1x DPBS (Dulbeccos’s, with 4 mM MgCl_2_, pH 7.3), followed by incubation with 1 mM ortho- phenyldimaleimide (o-PDM) for 30-60 minutes. A 12 μl sample with membranes was also diluted into 700 μl of 1x DPBS, followed by incubation with 1 mM o-PDM for 10-15 minutes on ice before 70 μl of 10% C_12_M was added, and incubation was continued for another 30-60 minutes. The PDM reaction was terminated by adding 50 μl of DTT and incubated at room temperature for 5 min and on ice for 10 minutes. Then, the reaction was centrifuged through a sucrose cushion (120 μl; 0.5 M sucrose, 100 mM KCl, 5 mM MgCl_2_, 50 mM HEPES (pH 7.3) and 1mM DTT) at 4°C for 22 min at 70,000 rpm using a TLA 100.3 Beckman Optima TLX Ultracentrifuge, to isolate ribosome-bound peptide. The pellets were resuspended in 40 μl of PBS 2% SDS and 2 mM DTT at RT for 30 minutes. Samples were diluted with 60 μl DPBS buffer containing 10 mM methoxy- polyethylene glycol maleimide (PEG-MAL, MW 5000; SunBio Korea) for a final concentration of 6 mM and incubated at RT for 2 hrs. After pegylation, samples were precipitated with acetone containing 0.4 mM HCl overnight. The supernatant was aspirated and the pellet resuspended in 14.5 μl DPBS (1x), 2.5 μl of DTT (10x) and 7 μl of LDS (4x). Samples with truncated constructs were treated with 50 μg/ml RNase and incubated at room temperature for 20 min. Intramolecular *Pxlink* was calculated as described in Tu et al.^13^

### Gel electrophoresis and fluorography

All final samples were heated at 70 °C for 10 min in 1× of NuPAGE loading buffer (Invitrogen, Waltham, MA) before loading onto the NuPAGE gel (Invitrogen). Electrophoresis was performed using the NuPAGE system and precast Bis-Tris 10 %, 12 %, or 4–12 % gradient gels and MES or MOPS running buffer. Gels were soaked in Amplify (Amersham, ArlingtonHeights, IL) or Enlightning (Revvity Health Sciences, Boston, MA) to enhance ^35^S fluorography, dried and exposed to Hyperfilm ECL film at −80 °C. Typical exposure times were 16–30 h. Quantification of gels was carried out directly using a Molecular Dynamics PhosphorImager (Typhoon FLA 9500) and ImageQuant TL (v7.0) (Sunnyvale, CA).

## ACKNOWLEDGEMENTS

This work was supported by NIH grant GM52302 to C.D.

## REFERENCES

1. Shishido, H., Yoon, J. S., Yang, Z. & Skach, W. R. CFTR trafficking mutations disrupt cotranslational protein folding by targeting biosynthetic intermediates. Nat.Commun. 11, 4258 (2020).

2. Sander, I. M., Chaney, J. L. & Clark, P. L. Expanding Anfinsen’s principle: contributions of synonymous codon selection to rational protein design. J.Am.Chem.Soc. 136, 858–861 (2014).

3. Anderson, C. L. et al. Large-scale mutational analysis of Kv11.1 reveals molecular insights into type 2 long QT syndrome. Nat.Commun. 5, 5535 (2014).

4. Anderson, C. L. et al. Most LQT2 mutations reduce Kv11.1 (hERG) current by a class 2 (trafficking-deficient) mechanism. Circulation 113, 365–373 (2006).

5. Smith, J. L. et al. Molecular pathogenesis of long QT syndrome type 2. J.Arrhythm. 32, 373–380 (2016).

6. Lu, J., Robinson, J. M., Edwards, D. & Deutsch, C. T1-T1 interactions occur in ER membranes while nascent Kv peptides are still attached to ribosomes. Biochemistry 40, 10934–10946 (2001).

7. Kosolapov, A. & Deutsch, C. Folding of the voltage-gated K+ channel T1 recognition domain. Journal of Biological Chemistry 278, 4305–4313 (2003).

8. Kosolapov, A., Tu, L., Wang, J. & Deutsch, C. Structure Acquisition of the T1 Domain of Kv1.3 During Biogenesis. Neuron 44, 295–307 (2004).

9. Robinson, J. M. & Deutsch, C. Coupled Tertiary Folding and Oligomerization of the T1 Domain of Kv Channels. Neuron 45, 223–232 (2005).

10. Lu, J. & Deutsch, C. Folding zones inside the ribosomal exit tunnel. Nat.Struct.Mol.Biol. 12, 1123–1129 (2005).

11. Kosolapov, A. & Deutsch, C. Tertiary interactions within the ribosomal exit tunnel. Nat.Struct.Mol.Biol. 16, 405–411 (2009).

12. Tu, L. W. & Deutsch, C. A folding zone in the ribosomal exit tunnel for Kv1.3 helix formation. J.Mol.Biol. 396, 1346–1360 (2010).

13. Tu, L., Khanna, P. & Deutsch, C. Transmembrane segments form tertiary hairpins in the folding vestibule of the ribosome. J.Mol.Biol. 426, 185–198 (2014).

14. Tu, L. & Deutsch, C. Determinants of Helix Formation for a Kv1.3 Transmembrane Segment inside the Ribosome Exit Tunnel. J.Mol.Biol. 429, 1722–1732 (2017).

15. Li, M. J. Y. N. a. J. L. Y. Specification of subunit assembly by the hydrophilic amino-terminal domain of the Shaker potassium channel. Science 257, 1225–1230 (1992).

16. Shen, N. V., Chen, X., Boyer, M. M. & Pfaffinger, P. Deletion analysis of K+ channel assembly. Neuron 11, 67–76 (1993).

17. Zerangue, N., Jan, Y. N. & Jan, L. Y. An artificial tetramerization domain restores efficient assembly of functional Shaker channels lacking T1. Proceedings of the National Academy of Sciences of the United States of America 97, 3591–3595 (2000).

18. Shi, G. et al. Beta subunits promote K+ channel surface expression through effects early in biosynthesis. Neuron 16, 843–852 (1996).

19. Yu, W., Xu, J. & Li, M. NAB domain is essential for the subunit assembly of both α-α and α-ß complexes of Shaker -like potassium channels. Neuron 16, 441–453 (1996).

20. Gulbis, J. M., Zhou, M., Mann, S. & MacKinnon, R. Structure of the cytoplasmic beta subunit-T1 assembly of voltage-dependent K+ channels. Science 289, 123–127 (2000).

21. Deutsch, C. Potassium channel ontogeny. Annual.Review of Physiology. 64, 19–46 (2002).

22. Deutsch, C. The birth of a channel. Neuron 40, 265–276 (2003).

23. Gu, C., Jan, Y. N. & Jan, L. Y. A conserved domain in axonal targeting of Kv1 (Shaker) voltage-gated potassium channels. Science 301, 646–649 (2003).

24. Cushman, S. J. et al. Voltage dependent activation of potassium channels is coupled to T1 domain structure. Nature Structural Biology 7, 403–407 (2000).

25. Minor, D. L. et al. The polar T1 interface is linked to conformational changes that open the voltage-gated potassium channel. Cell 102, 657–670 (2000).

26. Kurata, H. T. et al. Amino-terminal determinants of U-type inactivation of voltage-gated K+ channels. Journal of Biological Chemistry 277, 29045–29053 (2002).

27. Wang, G. & Covarrubias, M. Voltage-dependent gating rearrangements in the intracellular T1-T1 interface of a K+ channel. J.Gen.Physiol 127, 391–400 (2006).

28. Matteson, D. R. & Deutsch, C. K channels in T lymphocytes: a patch clamp study using monoclonal antibody adhesion. Nature 307, 468–471 (1984).

29. Price, M., Lee, S. C. & Deutsch, C. Charybdotoxin inhibits proliferation and interleukin 2 production in human peripheral blood lymphocytes. Proc.Natl.Acad.Sci.U.S.A. 86, 10171–10175 (1989).

30. DeCoursey, T. E., Chandy, K. G., Gupta, S. & Cahalan, M. D. Voltage-gated K+ channels in human T lymphocytes: a role in mitogenesis? Nature 307, 465–468 (1984).

31. Chandy, K. G., DeCoursey, T. E., Cahalan, M. D., McLaughlin, C. & Gupta, S. Voltage-gated potassium channels are required for human T lymphocyte activation. Journal of Experimental Medicine 160, 369–385 (1984).

32. Deutsch, C. K+ channels and mitogenesis. Prog.Clin.Biol.Res. 334, 251–271 (1990).

33. Serrano-Albarras, A., Estadella, I., Cirera-Rocosa, S., Navarro-Perez, M. & Felipe, A. Kv1.3: a multifunctional channel with many pathological implications. Expert.Opin.Ther.Targets. 22, 101–105 (2018).

34. Huang, Z., Hoffman, C. A., Chelette, B. M., Thiebaud, N. & Fadool, D. A. Elevated Anxiety and Impaired Attention in Super-Smeller, Kv1.3 Knockout Mice. Front Behav.Neurosci. 12, 49 (2018).

35. Xu, J. et al. The voltage-gated potassium channel Kv1.3 regulates energy homeostasis and body weight. Hum.Mol.Genet. 12, 551–559 (2003).

36. Long, S. B., Tao, X., Campbell, E. B. & MacKinnon, R. Atomic structure of a voltage-dependent K+ channel in a lipid membrane-like environment. Nature 450, 376–382 (2007).

37. Selvakumar, P. et al. Structures of the T cell potassium channel Kv1.3 with immunoglobulin modulators. Nat Commun 13, 3854 (2022). 10.1038/s41467-022-31285-5

38. Tu, L., Wang, J. & Deutsch, C. Biogenesis of the T1-S1 linker of voltage-gated K+ channels. Biochemistry 46, 8075–8084 (2007).

39. Munoz, V. & Serrano, L. Development of the multiple sequence approximation within the AGADIR model of alpha-helix formation: comparison with Zimm-Bragg and Lifson-Roig formalisms. Biopolymers 41, 495–509 (1997).

40. Cai, Y. C., Osborne, P. B., North, R. A., Dooley, D. C. & Douglass, J. Characterization and functional expression of genomic DNA encoding the human lymphocyte type n potassium channel. DNA.Cell.Biol. 11, 163–172 (1992).

41. Hopkins, W. F., Demas, V. & Tempel, B. L. Both N- and C-terminal regions contribute to the assembly and functional expression of homo- and heteromultimeric voltage-gated K+ channels. J.Neurosci. 14, 1385–1393 (1994).

42. Kobertz, W. R. & Miller, C. K+ channels lacking the ‘tetramerization’ domain: implications for pore structure. Nature Structural Biology 6, 1122–1125 (1999).

43. Tu, L. et al. Voltage-gated K + Channels Contain Multiple Intersubunit Association Sites. Journal of Biological Chemistry 271, 18904–18911 (1996).

44. Sheng, Z., Skach, W., Santarelli, V. & Deutsch, C. Evidence for Interaction between Transmembrane Segments in Assembly of Kv1.3. Biochemistry 36, 15501–15513 (1997).

45. Lee, T. E., Phillipson, L. H., Kuznetsov, A. & Nelson, D. J. Structural determinant for assembly of mammalian K+ channels. Biophysical Journal 66, 667–673 (1994).

46. Kobertz, W. R., Williams, C. & Miller, C. Hanging gondola structure of the T1 domain in a voltage-gated K(+) channel. Biochemistry 39, 10347–10352 (2000).

47. Lu, J. & Deutsch, C. Pegylation: a method for assessing topological accessibilities in Kv1.3. Biochemistry 40, 13288–13301 (2001).

48. Capera, J. et al. A novel mitochondrial Kv1.3-caveolin axis controls cell survival and apoptosis. Elife 10 (2021). 10.7554/eLife.69099

49. Lehmann, E. L. & Casella, G. Theory of Point Estimation. (Springer-Verlag, 1998).

50. Tu, L. & Deutsch, C. Evidence for dimerization of dimers in K+ channel assembly. Biophysical Journal 76, 2004–2017 (1999).

51. Tu, L., Santarelli, V. & Deutsch, C. Truncated K+ channel DNA sequences specifically suppress lymphocyte K+ channel gene expression. Biophysical Journal 68, 147–156 (1995).

52. Strang, C., Cushman, S. J., DeRubeis, D., Peterson, D. & Pfaffinger, P. J. A central role for the T1 domain in voltage-gated potassium channel formation and function. Journal of Biological Chemistry 276, 28493–28502 (2001).

53. Van Dongen, A. M. J., Frech, G. C., Drewe, J. A., Joho, R. H. & Brown, A. M. Alteration and deletion of K+ channel function by deletions at the N- and C-temini. Neuron 5, 433–443 (1990).

54. Aiyar, J., Grissmer, S. & Chandy, K. G. Full-length and truncated Kv1.3 K+ channels are modulated by 5-HTic receptor activation and independently by PKC. Am.J.Physiol. 265, C1571–C1578 (1993).

